# Phylogenetic and structural analyses reveal the determinants of DNA binding specificities of nucleoid-associated proteins HU and IHF

**DOI:** 10.1101/057489

**Authors:** Debayan Dey, Valakunja Nagaraja, Suryanarayanarao Ramakumar

**Author notes:** **Corresponding author** Suryanarayanarao Ramakumar.

## Abstract

Nucleoid-associated proteins (NAPs) are chromosome-organizing factors, which affect the transcriptional landscape of a bacterial cell. HU is an NAP, which binds to DNA with a broad specificity while homologous IHF (Integration Host Factor), binds DNA with moderately higher specificity. Specificity and differential binding affinity of HU/IHF proteins towards their target binding sites play a crucial role in their regulatory dynamics. Decades of biochemical and genomic studies have been carried out for HU and IHF like proteins. Yet, questions related to their DNA binding specificity, and differential ability to <u>bend</u> DNA thus affecting the binding site length remained unanswered. In addition, the problem has not been investigated from an evolutionary perspective. Our phylogenetic analysis revealed three major clades belonging to HU, IHFα and IHFβ like proteins with reference to *E. coli*. We carried out a comparative analysis of three-dimensional structures of HU/IHF proteins to gain insight into the structural basis of clade division. The present study revealed three major features which contribute to differential DNA binding specificity of HU/IHF proteins, I) conformational restriction of DNA binding residues due to salt-bridge formation II) the enrichment of alanine in the DNA binding site increasing conformational space of flexible side chains in its vicinity and III) nature of DNA binding residue (Arg to Lys bias in different clades) which interacts differentially to DNA bases. Differences in the dimer stabilization strategies between HU and IHF were also observed. Our analysis reveals a comprehensive evolutionary picture, which rationalizes the origin of multi-specificity of HU/IHF proteins using sequence and structure-based determinants, which could also be applied to understand differences in binding specificities of other nucleic acid binding proteins.

## Introduction

Nucleoid-associated proteins (NAPs) play a crucial architectural role in DNA bending and compaction as well as a regulatory role in various DNA transaction processes like replication and recombination (Dorman, 2009; Dillon and Dorman, 2010). In recent years, there has been an increasing interest in exploring the structure as well as the gene regulatory network involving NAPs (Browing et al., 2010; Prieto et al., 2012; Tolstorukov et al., 2016). They are one of the most abundant proteins in bacteria and exhibit promiscuity in DNA binding, affecting transcription of many genes.

HU is a dimeric NAP and a major component of bacterial nucleoid (Rouviere-Yaniv and Gros, 1975). HU and Integration Host Factor (IHF) belong to prokaryotic DNA-bending protein family (SCOP classification) and consist of three alpha helices and five beta strands, where the beta strands from each protomer form the DNA binding cradle while the alpha helices form the dimerization core representing the HU/IHF fold. They are also present in archaea, viruses and chloroplasts of many eukaryotes (Borca et al., 1996, Kobayashi et al., 2002, White and Bell, 2002). HU shares both structural and sequence similarity to Integration host factor (IHF), a homologous protein which is more sequence-specific in its DNA binding than HU. HU and IHF play crucial architectural roles in bacterial DNA condensation and additionally play a regulatory role in many cellular processes. They are involved in replication by binding to OriC region (Ryan et al., 2002), DNA recombination and repair (Kamashev and Rouviere-Yaniv, 2000), cell division (Dri et al., 1991) and functional interaction with DNA supercoiling maintaining proteins like gyrases and topoisomerases (Bensaid et al., 1996; Malik et al., 1996). HU/IHF proteins play the role of both repressor and activator for various genes in a well-orchestrated spatiotemporal manner (Aki et al., 1996; Oborto et al., 2008). HU facilitates bending of the DNA to bring the two distal GalR binding sites near, which helps in the formation of tetrameric GalR complex (Kar and Adhya, 2001). Similarly, in site-specific DNA inversion by Hin recombinase, HU plays a role in assembly of invertasome (Haykinson and Johnson 1993). *E. coli* IHF also plays role in the functioning of CRISPR-Cas mediated adaptive immunity against foreign genetic elements by inducing sharp DNA bend at the AT rich leader sequence in the CRISPR loci, allowing the Cas1-Cas2 integrase to catalyze the integration *in vivo* (Nuñez et al.,2016). A direct regulation of Topoisomerase I by *Mycobacterium tuberculosis* HU (MtbHU) was recently reported, to enhance its relaxation activity (Ghosh et al., 2015). MtbHU is also acetylated at the C-terminal lysine enriched region, similar to eukaryotic histones and interacts with acetyl transferase Eis, which causes reduced DNA interaction and alteration in DNA compaction (Ghosh et al., 2016). HU also influences the host immune response, making it one of the key components in host-pathogen interaction (Kunisch et al., 2012). Moreover, in many organisms HU/IHF proteins are essential, which makes them a promising drug target for infectious pathogens. In view of their importance, in an earlier study, our group has determined the structure of MtbHU, which is essential for the organism and inhibited it using structure-based designed stilbene derivatives (Bhowmick et al., 2014).

Although sharing structural similarity, the DNA binding and bending features of HU and IHF are strikingly different, allowing them to selectively regulate genes from different genomic locations (Prieto et al., 2012). HU binds to DNA in a sequence promiscuous (which may be called as multi-specificity) manner while IHF is moderately sequence specific (Bonnefoy and Rouviere-Yaniv, 1991; Swinger and Rice, 2004). *E. coli* HU binds to random DNA sequence with a K_d_ of 200 - 2500 nM, whereas *E. coli* IHF binds to such sequences 100 times more weakly (K_d_=20 - 30mM). However, *E. coli* IHF strongly binds (K_d_=2-20 nM) to its cognate recognition sites (Swinger and Rice, 2004). Other differences between HU and IHF are related to binding site length and DNA bending angle (Swinger and Rice, 2004). The molecular mechanism of DNA binding multi-specificity (differential specificity with varied binding affinity) of HU/IHF proteins remains enigmatic, as little attention has been paid to the determinants at the sequence level. Hence, we asked the question of how the differential specificities can be rationalized in a structural and evolutionary framework.

To understand the sequence determinants, which influence the degree of DNA binding specificity, we undertook a phylogenetic study in conjunction with analysis of threedimensional structures. Our phylogenetic analysis revealed three major clades, belonging to HU, IHFα, and IHFβ like proteins with reference to *E. coli*. We noted a set of positions, which discriminates between HU and IHF clade proteins. We observed statistically significant amino acid compositional bias in the DNA binding sites of HU and IHF clade proteins and between different HU proteins from *Proteobacteria, Firmicutes*, and *Actinobacteria*. Our analysis rationalizes the molecular determinants of binding and bending differences in *E. coli* HU and IHF, one of the model systems for understanding differential binding. We also elucidate the differential dimer stabilization strategies in HU and IHFs, which might influence DNA binding and bending. We propose that the molecular mechanisms giving rise to specificity or multi-specificity depends on a combinatorial effect of the amino acid composition of the binding site, its flexibility, ionic and steric constraints.

The determinants themselves are more general in nature and the methodology from the present report can be applied in explaining multi-specificity in other nucleic acid binding proteins.

## Results and discussion

The determinants that influence the DNA binding specificities in HU and IHF like proteins remained an unresolved issue for long time. Earlier studies showed that *E. coli* HU binds DNA with moderate affinity but poor selectivity (rather mostly non-specific), while *E. coli* IHF binds to its cognate sequence with higher affinity (Swinger and Rice 2004). The questions addressed here are i) The molecular determinants guiding the evolution of paralogous proteins HU and IHFs, with varied DNA binding specificity ii) Can we gain some insight into the DNA binding specificities of other HU homologs from *Firmicutes* and *Actinobacteria* phyla, belonging to various pathogens e.g. *Mycobacterium tuberculosis, Staphylococcus aureus* etc. from the above determinants.

In the past, evolutionary trace (ET) method has been widely used to identify protein functional determinants and to find class specific residues (Innis et al., 2000; Sowa et al., 2001; Chakravarty et al., 2005). The evolutionary trace (ET) method that uses phylogenetic tree-based sequence alignments along with structural information has been successfully utilized to detect functional sites in many different protein families. In this method, a trace is generated by comparing the consensus sequences for a group of proteins, which originate from a common node in a phylogenetic tree and characterized by a common evolutionary time cut-off (ETC) (Innis et al., 2000). It classifies the residue positions as i) conserved across the family, ii) clade specific or iii) neutral. In comparison to other phylogenetic methods, its strength lies in its flexibility, which allows for a wide range of `functional resolutions’ (Innis et al., 2000).We have used ET method here to understand the phylogenetic grouping of HU/IHF proteins as well to understand here the clade specific residues. To further validate the clade division of HU/IHF family proteins, we created a Maximum Likelihood (ML) phylogenetic tree, which concurred with our ET results.

In our present analysis, we further utilized the ET method to consider the conservation and variability based on class specific residue groups (grouping based on physicochemical properties of amino acids) rather than the amino acids themselves. Specificity determining factors cannot always be absolutely conserved and mutate among amino acids with similar physicochemical properties, thus, scoring their positional conservation based on physicochemical properties seemed a rational choice. To understand the conservation and variability among different HU and IHF clades, we determined and compared class specific Shannon’s entropy to find discriminating positions. We also utilized hydrophobicity scale and pI of each sequence position to discriminate HU and IHF further. Unlike most of the enzymes where only a few residues (positions) are responsible for substrate selectivity and catalysis, HU/IHF proteins bind to DNA with a large interface spanning more than ~30 residues. Thus, selectivity might not always be dependent on just specific positions, rather on the amino acid composition of the entire DNA binding region. Thus, we performed an analysis, which can bring out the enrichment and depletion patterns of certain amino acids or groups between HU and IHF clades in a statistically significant manner. Previously, studies have shown subfamily (clade) specific amino acid compositional bias could be used for their classification (Bhasin and Raghava, 2004). In this report, we utilized the above stated phylogenetic, entropy based conservation and amino acid compositional analysis along with structural studies on DNA bound and apo proteins, to rationalize the determinants of specificity in HU and IHF. We also performed the same analysis to determine differences in HU clade members of different phyla.

## Taxonomic distribution and phylogenetic analysis reveal three clades of HU/IHF homologs spread across bacteria

A diverse set of sequences homologous to HU/IHF fold was found ubiquitously distributed among bacteria through systematic database searches (Supplementary Table S1). In *Proteobacteria* and *Bacteroidetes*, they are present in multiple paralogous copies, while in other phyla, it is mostly present as a single copy. In *Streptomyces* genus, it has two paralogs, one similar to *E. coli* HU which is expressed in growth phase while the other contains an additional C-terminal extension (~ 100 residues), enriched in lysine and alanine, expressed during the sporulation phase (Salerno et al., 2009). These positively charged tails, which are also present in *Deinococcus radiodurans* HU (DrHU) precede the HU/IHF fold and form a 47-amino acid long extension, which binds to DNA (Ghosh and Grove, 2006). We noted that HU homologs of *Mycobacterium smegmatis* and *Mycobacterium tuberculosis* along with many other in *Actinobacteria* are also associated with a similar lysine rich “PAKKA” repeat in the C-terminal, were it is implicated in protection of DNA from adverse conditions. This study found several *Betaproteobacterial* HU with N-terminal positively charged tails associated with HU-IHF fold, which was not reported earlier (Supplementary Table 2). We also observed that viral genomes of African swine fever virus and *Staphylococcus phage* code for HUlike proteins, which have substantial similarity to *Proteobacterial* HU and might have originated from them. While the role of bacteriophage coded HU homologs in host integration is well known, its role in eukaryotic viruses’ is not understood. HU like proteins are also present in eukaryotic organisms with apicoplast, which shares sequence similarity to endosymbiont nitrogen fixing *Rhizobial* HU. In eukaryotic pathogens, *Plasmodium* and *Toxoplasma* plastids contain circular DNA that is organized by HU homologs and are essential for the organisms’ survival (Ram et al., 2008).

We performed phylogenetic studies on all HU/IHF proteins from bacteria, with the exception of phylum *Bacteroidetes* due to its highly divergent sequences which resulted in poor multiple sequence alignment (MSA). The alignment of the dataset with all bacteria is termed as MSA1 from here on. The evolutionary trace (ET) phylogenetic tree was partitioned into 20 traces (Fig. 1) with first few traces (Trace 1-4) dividing it into amajor clade (86% sequences) and other smaller clades (14% sequences). The 5^th^ trace divided the tree into three major clades, along with many smaller ones. Further traces divided the HU and IHF clades into smaller sub-groups, which is not of this studies interest. The three major clades belong to a) HU like proteins, with distribution throughout bacteria and b) IHFα and c) IHFβ like proteins (with reference to *E. coli* IHF) from mostly *Proteobacteria*. Although IHFα and IHFβ are mostly restricted to *Proteobacteria*, their average identities within the clades were found to be 42% and 41% respectively, while the average identity of HU clade was found to be 39% in our dataset. The clade divisions made by a maximum likelihood (ML) tree concurred with the results of the ET phylogenetic tree (Supplementary Fig. S1).

**Figure 1.**
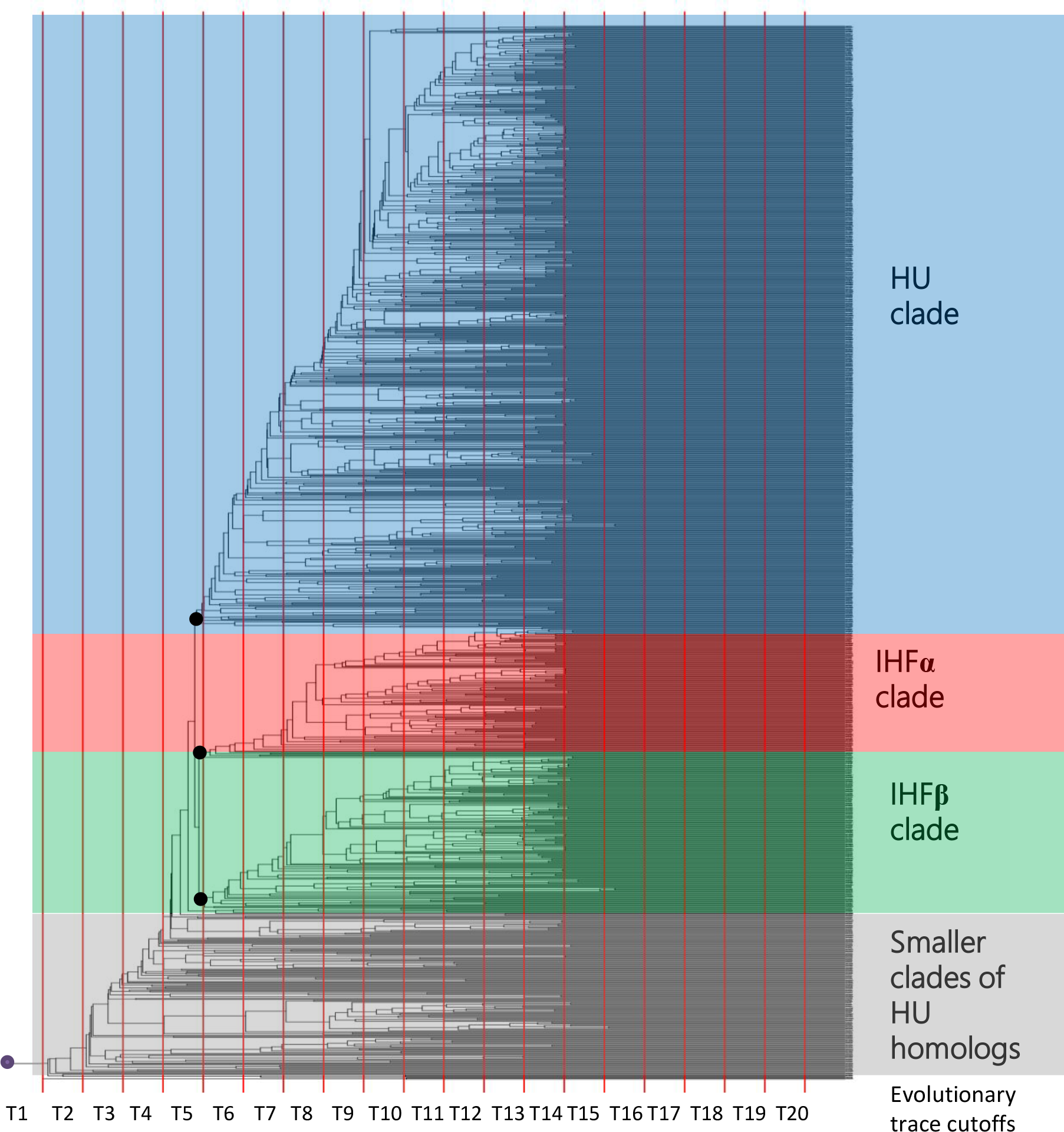
Evolutionary trace Phylogenetic tree. Showing three major clades namely, HU clade (red color shaded), IHFα (blue color shaded) and IHFβ (green color shaded) clades. Evolutionary trace cutoff (ETC) are shown as T1-T20 and the clade division takes place at 5^th^ trace (shown by black nodes).

As *E. coli* IHF (heterodimer formed from IHFα and IHFβ chain) and *E. coli* HU are taken as model systems to understand different DNA interaction features in HU/IHF family proteins in the past, we speculated that the DNA binding features like bendability and specificity of various other HU homologs might be different across phyla. Also, our analysis found that other than *Proteobacteria* and *Bacteroidetes*, mostly all other phyla contains a single copy of HU like proteins, which has to perform comparable functional role of both HU and IHF. Thus, their nature must be slightly different from *Proteobacterial* HU which would have imprinted during their sequence evolution.

To understand the phyla-specific variations in HU homologs affecting the differential specificity, we created three different sequence alignments of a) *Proteobacterial*, b) *Firmicutes* and c) *Actinobacterial* HU/IHF homologs and performed phylogenetic analysis using ET method.

Our analysis found *Proteobacterial* HU/IHF homologs were branched into three major clades at 6^th^ trace consistent with the previous all bacterial dataset of HU/IHF homologs (MSA1 dataset). *Actinobacterial* HU homologs were divided into two clades at 2^nd^ trace, where the first clade (M1) consists of HU from *Mycobacterium tuberculosis*, associated with C-terminal Lys and Ala enriched motifs (more than >90% sequences). The second clade (M2) consists mostly of HU like proteins which don’t have any positive charged N-or C-terminal extensions. Although only the HU/IHF fold was considered in this study, yet, the MSA of these proteins could segregate them based on their domain arrangements e.g. the presence of C-terminal extensions. Thus, we speculate that the presence of additional C-terminal region must have an evolutionary imprint on its HU/IHF fold. Similarly, our analysis of *Firmicutes* HU homologs found no large clade divisions in traces (1-5) and the whole dataset is taken for further analysis.

## Evolutionary trace analysis identifies invariant and class-specific residues in HU and IHF

Proteins are composed of a set of residues which are conserved either to maintain its fold but also evolves a set of residues which influences specificity (class specificity in terms of recognizing specific binding partners). We divided the protein into two segments, i) β-sheet DNA binding region (BDR) and ii) alpha helical region (AHR) (Fig. 2a,b). To understand the invariant and class-specific residues in HU/IHF fold proteins, we analyzed the MSA of HU, IHFα, and IHFβ at 5^th^ traces separately to decipher the positional determinants which can discriminate HU from IHFs. Normalized Shannon’s entropy of MSA1 shows the residue conservation and variability in different clades of HU/IHF proteins (Fig.2c), in which we observed that the residues involved in DNA binding region are more conserved than the residues in dimerization region. In the present report, we have used MtbHU sequence as representative while describing sequence positions in the alignment.

**Figure 2.**
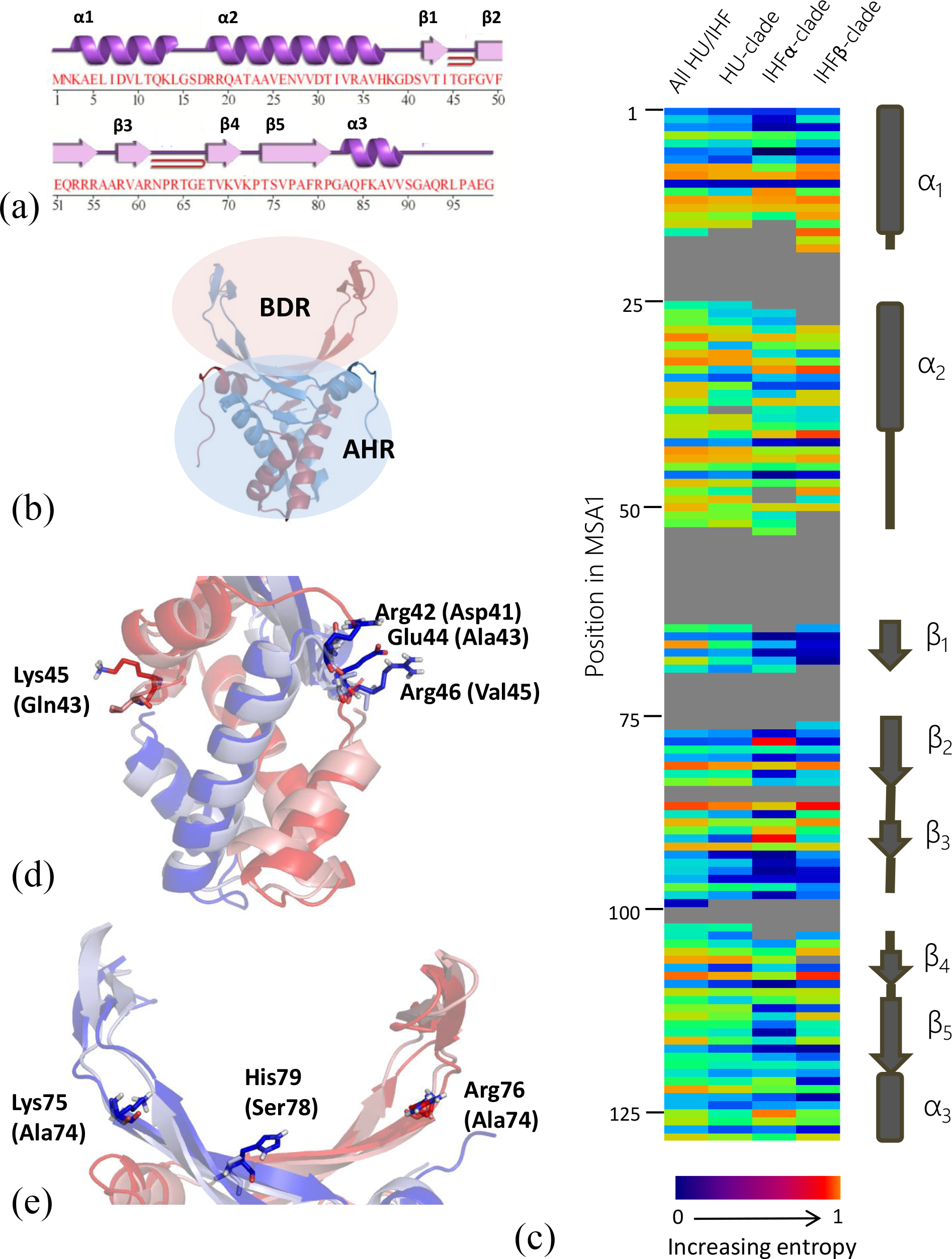
Conservation and variability among clades. **(a)** MtbHU^N^ (1-99) sequence with secondary structural elements. **(b)** The structure of MtbHU^N^(PDB ID: 4PT4) which is determined at our laboratory is shown is cartoon representation. The alpha helices α_1_,α_2_ and α_3_ (termed as alpha helical region or AHR) from both protomers form the helical bundle for dimerization while β_1_-β_5_ form the DNA binding cradle (termed as beta sheet DNA binding region or BDR). **(c)**Heat map ofnormalized Shannon’s entropy for the following alignments: Complete alignment of all bacterial HU-IHF dataset of 1112 sequences (MSA1) (lane 1), HU clade (lane 2), IHFα (lane 3) and IHFβ (lane 4). The data shows alpha helical region (1-50 and 120-128 in alignment) is moderately conserved, while beta strand region (51-119 in alignment) is more conserved. **(d)** IHFα clade specific residue Lys45 is replaced by Glu43 in HU (shown in the structural alignment of *E. coli* IHF and Modeled *E. coli* HUαβ, represented as red for α-chains and blue for β-chains, the bracketed residues are from *E. coli* HU), while in IHFβ, Arg42- Arg 46 is replaced by Asp41-Val45, respectively. Glu44 replaces the Ala43 in IHFβ clade. Thus, these crucial positively charged residues in the DNA draping site is exclusive to IHF clades and are responsible for stabilizing the bent DNA and hence increasing the binding site length. **(e)** In the DNA binding site, IHF clades has some exclusive positively charged residues like Lys75 (in IHFβ) and Arg76 (in IHFα), along with aromatic His residue His79 (in IHFβ), which are replaced by non-charged residues in HU.

We observed that the hydrophobic residues in the AHR (Supplementary Table S3) which are primarily responsible for the formation of dimerization core are conserved in the entire alignment. The BDR also consists of some highly invariant residues which form the aromatic core, such as F47, F50, and F79. Other invariant residues include G46 and P63, alongside hydrophobic amino acids in BDR (alignment position 66, 67, 107 and 109 in MSA1). P63 is the residue involved in DNA intercalation and generates hinges, which are further stabilized by neighboring residues (Chen et al., 2004). R53 of MtbHU (position 84 in MSA1) is moderately conserved while R55 and R58 are highly conserved positions (Fig. 2c). While R55 interacts with the backbone phosphates of the DNA in most of the HU/IHF proteins, R58 interacts with its DNA base. In HU clade, Arg occupies position 88 corresponding to R55 in MtbHU, while in IHFα clade, it is predominantly occupied by Lys. In IHFβ clade, it is moderately conserved, occupied by Arg, Tyr, and His. DNA binding residues R61 (near the conserved Pro tip), R80 and K86 are conserved throughout the HU/IHF proteins.

We further analyzed each position of HU, IHFα and IHFβ clades with respect to its Shannon’s entropy (SE), hydrophobicity and mean pI. Our analysis highlights various ways by which a residue can contribute to differential binding and DNA bending ability. Position 67 in MSA1 discriminates all the three clades, where in HU clade, it is moderately variable (SE=0.59), occupied by polar residues Thr, Ser and Gln.Whereas, in IHFα clade, it is occupied by Lys (K45 in *E. coli* IHFα) which interacts with DNA. In IHFβ clade, Glu (E44 in E.coli IHFβ) occupies it, which holds the two neighboring conserved DNA binding Arg (R42 and R46) (Fig. 2d). Similarly position 112 in MSA1, is variable in HU clade proteins, while is conserved in Arg (R76 in *E. coli* IHFα) in IHFα clade and Lys (K75 in *E. coli* IHFβ) in IHFβ clade, both of which interacts with the DNA (Fig. 2e). Furthermore, positions 87 and 116 in HU clade proteins are positively charged, while in IHFα clade dominantly negatively charged. Whereas, in IHFβ clade, these positions are enriched with His and aromatic residues, with *E. coli* IHFβ (H54 and H79) interacting with the DNA. Ser78 in *E. coli* HUβ is occupied by H79 in IHFβ(Fig. 2e). Other discriminatory position-specific differences in HU, IHFα, and IHFβ clade proteins are listed in Supplementary Table S4.

## Amino acid compositional differences in HU and IHF determines specificity

Charged amino acid residues on the surface of HU and IHF play a crucial role in determining its affinity and specificity for DNA binding. We observed significant (p-value <0.05) differences in charged (both positive and negative) and small amino acids’ (Gly, Ala, Pro, Ser) composition across HU and IHF clades (Table 1). Positively charged amino acids in the BDR are significantly enriched in IHF clades (23.5 *%* in IHFα and 25.7 % in IHFβ) compared to HU clade (20.0 %), while in the AHR, the percentage increase of positively charged residues in IHFβ is marginal compared to HU and IHFα. Keeping a similar trend, both BDR and AHR show a significant enrichment of negatively charged residues in IHF clades compared to HU. We noted an enrichment of Glu in the AHR of IHF clade proteins (10.2 % in IHFα and 10.6 % in IHFβ) compared to HU (7.6 %). We found that both positive and negatively charged residue enrichment and depletion is characteristic in different clades. Arg was significantly enriched in IHF clades (12.5 % in IHFα and 13.6 % in IHFβ) than HU (9.5 %) in the BDR and a similar trend is observed in the AHR too, where IHF (6.0 % in IHFα and 6.2 % in IHFβ) leads over HU (2.2 %). A reverse trend was observed for Lys, which shows significant depletion in the AHR of IHF clades (8.4 % in IHFα and 9.2 % in IHFβ) in comparison to HU (13.5 %). We also noted that the DNA phosphate and base interacting preferences are different for Arg and Lys, thus this bias can affect specificity (Luscombe et al. 2001). Arg interacts with DNA bases more often than Lys, thus making it a better choice as a specificity-improving residue than Lys. These differences in charged amino acids affect the salt-bridge making propensity, which influences the DNA binding specificity.

**Table 1.**
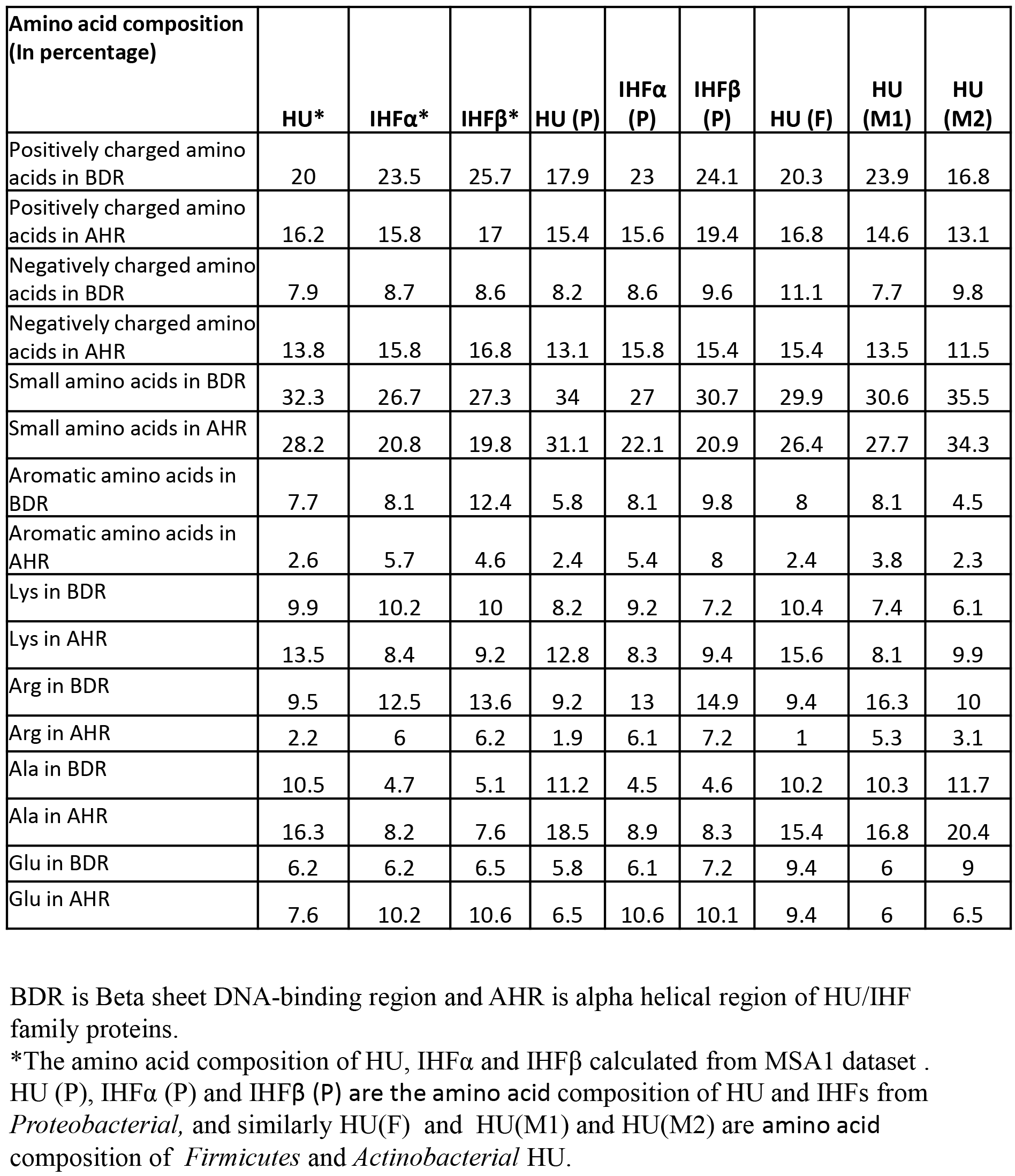
Comparison of amino acid compositional bias between HU and IHF clades and Phyla specific HU proteins.

## Salt bridges drive specificity by constraining flexibility in HU/IHF proteins

Earlier studies have hinted at the importance of salt bridge rearrangements at the DNA binding interface (Ma et al., 2010). A previously performed molecular dynamics study on *E. coli* IHF by Ma at al. showed K15, R21, R77 and K97 of IHFα and K69 of IHFβ have a high probability of forming salt bridges in native condition, but in DNA-bound structure (PDB: 1IHF) the salt bridges were missing (Ma et al., 2010). Due to IHFs enrichment of both positive and negatively charged residues in the BDR compared to HU, its salt-bridge making capacity is also much higher, as we observed from comparative structural analysis. Due to the absence of apo *E. coli* IHF structure and absence of the complete *E.coli* HUαβ structure (The beta sheet region is mostly disordered), a structural comparison was not possible, but from overall composition of charged residues of *E. coli* IHF vs. *E. coli* HUαβ gives a clear hint (Table 2). This trend is further observed in *Salmonella typhimurium coli* IHFαβ vs. HUαβ (Supplementary Table S5).

**Table 2.** Comparison of amino acid compositional bias between E.coli HU and IHF proteins.

| Amino acid composition<br>(In percentage) | HUα | HUβ | IHFα | IHFβ | HU($\alpha\beta$) | IHF($\alpha\beta$) |
| --- | --- | --- | --- | --- | --- | --- |
| Positively charged amino acids in BDR | 18.4 | 15.8 | 21.4 | 23.7 | 17.1 | 22.55 |
| Positively charged amino acids in AHR | 15.4 | 15.3 | 19.3 | 19.7 | 15.35 | 19.5 |
| Negatively charged amino acids in BDR | 7.9 | 10.6 | 11.9 | 13.1 | 9.25 | 12.5 |
| Negatively charged amino acids in AHR | 13.4 | 11.5 | 21.1 | 14.3 | 12.45 | 17.7 |
| Small amino acids in BDR | 29 | 34.2 | 23.8 | 29 | 31.6 | 26.4 |
| Small amino acids in AHR | 30.7 | 40.4 | 22.9 | 25.1 | 35.55 | 24 |
| Aromatic amino acids in BDR | 7.9 | 5.3 | 7.1 | 13.2 | 6.6 | 10.15 |
| Aromatic amino acids in AHR | 1.9 | 1.9 | 7.1 | 9 | 1.9 | 8.05 |
| Lys in BDR | 7.9 | 7.9 | 7.1 | 7.9 | 7.9 | 7.5 |
| Lys in AHR | 15.4 | 11.5 | 12.3 | 8.9 | 13.45 | 10.6 |
| Arg in BDR | 7.9 | 7.9 | 14.3 | 13.2 | 7.9 | 13.75 |
| Arg in AHR | 0 | 3.8 | 7 | 5.4 | 1.9 | 6.2 |
| Ala in BDR | 13.2 | 18.4 | 2.4 | 2.6 | 15.8 | 2.5 |
| Ala in AHR | 19.2 | 23.1 | 8.8 | 10.7 | 21.15 | 9.75 |
| Glu in BDR | 5.3 | 5.3 | 4.8 | 10.5 | 5.3 | 7.65 |
| Glu in AHR | 9.6 | 3.8 | 15.8 | 10.7 | 6.7 | 13.25 |

BDR is Beta sheet DNA-binding region and AHR is alpha helical region of HU/IHF family proteins.
HU( $\alpha\beta$ ) and IHF( $\alpha\beta$ ) are the average of both the chains which forms an functional dimer.

Although Firmicutes HUs belongs to HU clade, some of its amino acid compositional bias reflects those of IHFs (Table 1). Its negatively charged amino acid percentage is higher (11%) than its *Actinobacterial* (7.7% for M1 clade) or *Proteobacterial* (8.6%) counterpart (Table 1). A comparative analysis of crystal structures of *Staphylococcus aureus* HU (PDB: 4QJN and 4QJU) in native and DNA-bound form illustrate the concept of “making and breaking” of salt bridges (Kim et al., 2015). E51 in the native structure was interacting with K41 and K80 but it was additionally interacting with R53 in the DNA-bound form, which shows an alternate conformation (one interacting with DNA, while other making a salt bridge with E51), illustrating salt bridge dynamicity (Fig. 3a,b). Similarly, the salt bridge between E54 and K75 was disrupted from the native structure, and E54 formed a new salt bridge with R53. E54 displays different conformations in the same DNA-bound structure in different chains, which influences its interaction with R53 (Fig. 3c). Additionally, salt bridge rearrangement shifts the interaction of K59 from E68 and D70 (native) to only D70 (DNA bound) (Fig. 3d). In addition, a salt bridge between D87 and K90 is disrupted upon binding to DNA. Thus, salt-bridge forming capacity and its dynamicity both plays crucial role in the restricting the geometry of the DNA binding residues and thus affects specificity determination.

**Figure 3.**
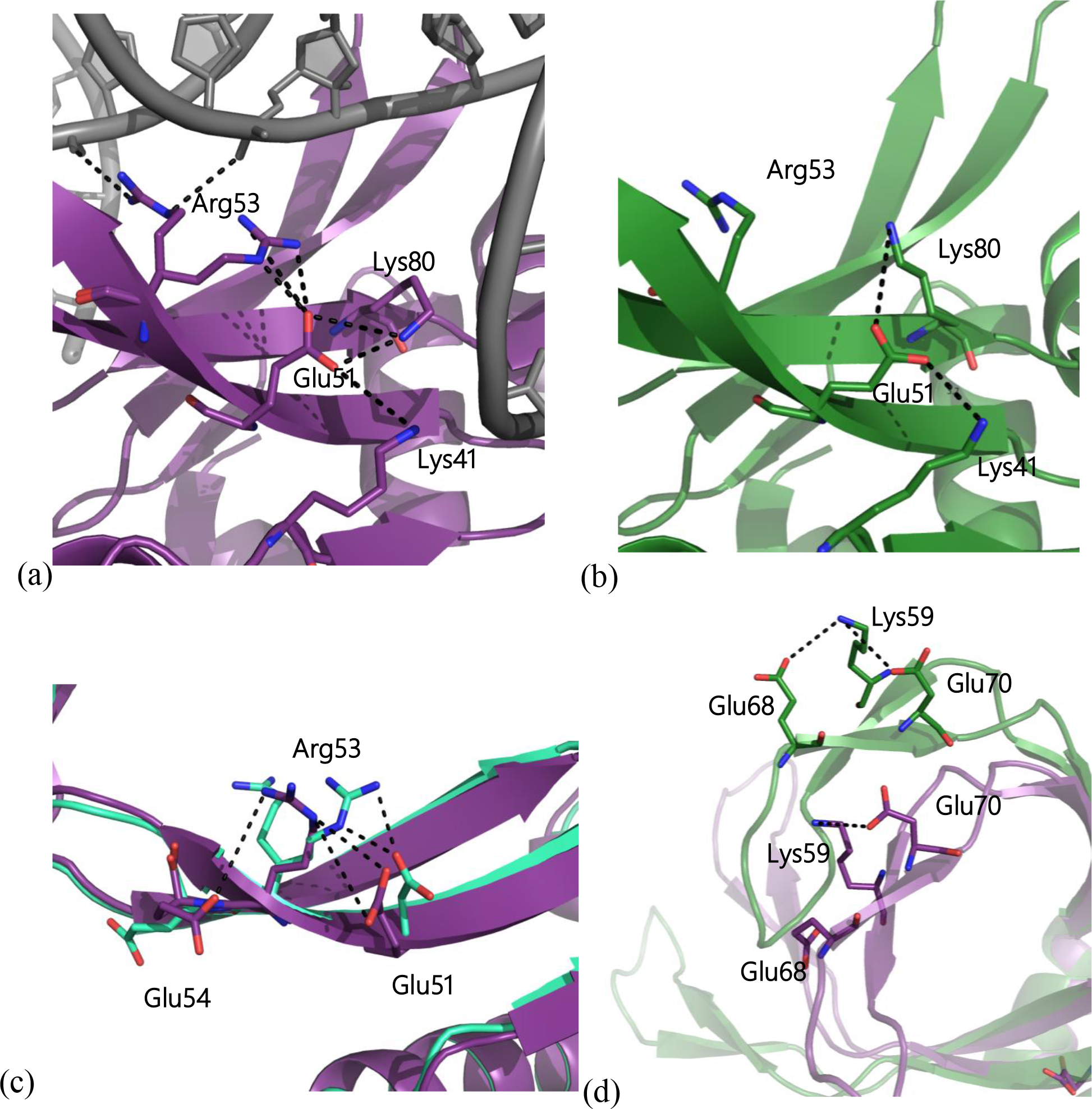
Salt bridge rearrangements and conformational flexibility promote promiscuity. **(a)** In DNA-bound form of HU in *Staphylococcus aureus*(PDB ID: 4QJU),Glu51 acts as a hub of electrostatic interactions, where it holds Lys41, Lys80 as well as Arg53, which in its alternate conformation binds to DNA. **(b)**In unbound form *Staphylococcus aureus*HU (4QJN), Glu51interacts with Lys41 and Lys80, but with different side chain conformer. This comparison gives an example of salt bridge rearrangements occurring at the DNA binding site which influences the promiscuity. **(c)**In *Staphylococcus aureus* HU (PDB ID: 4QJU) DNA-bound form, different chains in the dimer show salt bridge dynamicity. In chain B (Cyan-green) Arg 53 forms a salt bridge with Glu51, while in chain A, Arg 59 (violet) form salt bridge with Glu54. **(d)** In *Staphylococcus aureus* HU, unbound form (green),Lys59 form salt bridge with both Glu70 and Glu68, while in the DNA-boundform its interaction with Glu68 is lost.

The specificity and promiscuity of the protein towards its substrate is dependent on the overall flexibility of the binding residues or the active site region as a whole. Our results showing differences in specificity arising from differential salt-bridge forming capability, has been observed in other proteins too. Salt bridges, long-range electrostatic interactions, or even hydrogen bonding restricts this flexibility and thus promote specificity (Fornili et al., 2013). Our results are consistent with the previous findings of determinants of promiscuity in antibodies binding to hen egg-white lysozyme (HEL), which is dependent on the salt-bridge and other long-range electrostatic interactions, limiting flexibilities through geometric constraints(Sinha et al., 2002). They observed that promiscuous antibody (HH8) has the lowest number of Intra-and intermolecular hydrogen bonds and salt bridges, while the most specific antibody (HH26), shows an interweaving salt bridge network (Sinha et al., 2002). The above examples clearly validate our hypothesis that in HU/IHF proteins, amino acids compositional of positive and negatively charged residues contribute to salt bridge interactions and acts as a determinant of multi-specificity.

## Alanine in the DNA binding site drive multi-specificity by allowing flexibility

Continuing the argument of binding site flexibility driving promiscuity, small amino acid residues play a crucial role in rendering side chain flexibility of neighboring residues. In our present analysis, we observed a significant depletion of Ala (at both AHR and BDR) in IHF clades over HU (Table 1). HU clade proteins are enriched in Ala (10.5% in the BDR and 16.3 *%* in the AHR) compared to the IHF clade (6 *%* in IHFα and 6.2 *%* in IHFβ in the BDR; 4.7 % in IHFα and 5.1 % in IHFβ in the AHR) proteins in both regions. Ala provides an increase of conformational space for the rotatable side chains (of DNA binding residues) in its vicinity. The increased percentage of Ala in BDR thus increase the overall flexibility of the DNA binding site. Our results are justified by previous experiments on the effect of small residues in the neighborhood of DNA binding residues, which giving rise to promiscuous interactions in LEAFY family DNA binding protein (Sayou et al., 2014). Similar finding also support our results in which the flexibility of binding site in PDZ domain proteins’ are affected by small residues which results in its promiscuity (Münz et al., 2012). In contrast to small amino acid like Ala, we observed an enrichment of aromatic residues (in both AHR and BDR) for IHFβ clade proteins with 12.4% of aromatic residues in IHFβ compared to 7.7% in HU and 8.1% in IHFα in the BDR. In the AHR, both IHFα and IHFβ clade proteins are enriched in aromatic residues compared to HU, which results in a secondary aromatic cluster stabilizing the dimer interface (Table 1, discussed in later section).

## Multiple constraints discriminates the DNA binding specificities and bending angle of Proteobacterial HU and IHF

### Amino acid compositional differences

We took the *Proteobacterial* HU/IHF proteins as a model to study the various molecular determinants of differential specificities in these paralogs. HU proteins in *Proteobacteria* are significantly depleted in positively charged residues compared not only to *Proteobacterial* IHF clades but to also other HU proteins from *Actinobacteria* and *Firmicutes* (Table 1). Similarly, the small amino acid composition of both BDR and AHR are significantly enriched in *Proteobacterial* HU than IHFs and other HU clade proteins (Table 1). We also observed a depletion of Glu in both BDR (5.8% in HU, 6.1% IHFα and 7.2% IHFβ) and AHR (6.5% in HU, 10.6% IHFα and 10.1% in IHFβ) of *Proteobacterial* HU compared to IHFs (Table 1). In a similar trend, we observed an increase of both positive and negative charged residues in *E. coli* IHFα/β compared to HUα/β (Table 2). It also follows the previous trends of enrichment of small amino acid and depletion of aromatic amino acids in HUα/β compared to IHFα/β. We observed a Lys to Arg bias in the AHR where Lys is favored in HU while Arg is favored in IHF. In *Proteobacterial* HU, Arg constitutes only 1.9 % of the AHR in comparison to 6.1 % and 7.2 % in IHFα and IHFβ clades respectively. The enrichment of Arg in DNA binding BDR might facilitate its interaction with DNA base compared to Lys while in the AHR it serves a dual purpose of DNA binding and dimer stabilization by salt-bridge interactions.

### Exclusive salt bridges formed by IHF compared to HU influences specificity

The binding of IHF to DNA is coupled with the disruption, formation, and rearrangement of salt bridge interactions (Holbrook et al., 2001). Other than amino acid compositional differences in *Proteobacterial* HU and IHFs, we observed specific positions, which are exclusive to IHF compared to HU clade proteins. In the DNA binding cradle (BDR), positions corresponding to R76 and K88 of *E. coli* IHFα and H54, K75, and H79 of *E. coli* IHFβ are exclusive to IHF clade proteins. These extra positively charged residues, constrained by electrostatic and steric restrictions, impart specificity towards its DNA binding. One of the key salt bridge patterns observed exclusively in IHFβ clade is formed by R42, E44, and R46 in *E. coli* IHF β-chain (Supplementary Fig. S2). E44 is exclusive to IHFβ clade and holds R42 via a salt bridge, which in turn interacts with DNA. Our results are in agreement with earlier observations in which, IHF’s discrimination to its cognate binding site is affected upon the mutation of E44 to Ala (E44A). This mutation leads to disruption of the salt bridge and flexibility of R46, which can promiscuously bind to the mutated DNA site. Rice et al., 1996 argue that replacement of Glu at this position by other residues relaxes the specificity by allowing flexibility.

### Stabilizing interactions draw the DNA towards IHF than HU

At the site where DNA drapes down and interacts with the alpha helical region, IHF clade proteins has some exclusive residues K45 in IHF α and R42 and R46 in IHFβ, which stabilizes the bend. As the DNA drapes down, the presence of stabilizing interactions draw the DNA towards the alpha helical region (Khrapunov et al., 2006). We observed a stronger and more extended positively charged electrostatic potential along the draping path of IHF like proteins than HU (Supplementary Fig. S3a,b).

Even though positively charged residues are lined along the lateral side, their flexibility and accessibility are dependent their neighboring interactions. Although, earlier studies has shown that K3 forms a salt bridge with D26 in *Bacillus subtilis*, and its mutation to Ala (D26A) facilitates the conformational flexibility of K3 side chain by breaking the salt-bridge, and allowing it to engage with larger DNA, facilitating increased binding site length (Kamau et al., 2005), we observed this is not a general condition. Our study found, K3 is highly conserved across all the HU and IHF clades while E26 (position 35 in MSA1) is moderately conserved. In IHF clade a neighboring position (position 36 and 37 in MSA1) occupies a negatively charged residue with which K3 interacts via a salt bridge. The corresponding residues in *E. coli* IHF are K5 and E28 (IHFα), which interact via a salt bridge, while still interacting with DNA (Supplementary Fig. S3c). We observed a rearrangement of the salt bridge rather than its disruption. Now, it is understandable that the stabilization of DNA in the lateral section comes from a concerted effort of different positions in IHF mentioned earilier (K45 in IHFα and R42-R46 in IHFβ). Our analysis suggests that only stabilization of DNA by K3 position in not sufficient for stable bend angle and other positively charged residues in the DNA draping site may be required for extended site size and higher bending angle. Thus, we speculate that the bend angle and site size in various HU clade proteins are more dynamic than in IHF, and is dependent on salt concentration.

Therefore, a combinatorial effect of enrichment of smaller residues in the binding site providing conformational flexibility, and the presence of high salt bridge network in the DNA binding site accompanied by specific H-bonding render *Proteobacterial* IHF more specificity than HU. Similarly, due to the presence of exclusive positively charged residue in the DNA draping region and alternative salt-bridge rearrangements decide the DNA bent angle and binding site length.

### *Actinobacterial* and *Firmicutes* HU are different from their *Proteobacterial* counterpart

We observed a significant difference in percentage of charged amino acids (particularly positively charged) in the BDR of *Actinobacterial* (23.9% in M1 clade) and *Firmicutes* (20.3 %) HU compared to *Proteobacterial* HU (15.4%), while in the AHR, positively charged residue percentage is not significantly altered. We note that M2 group in *Actinobacteria* has fairly low positive charged residue percentage (16.8 % in the BDR) compared to any other HU clades. In *Firmicutes*, we observed a significant enrichment of negatively charged amino acids both in the BDR and AHR, compared to *Proteobacterial* or *Actinobacterial* HU (Table 1). Firmicutes HU consists 11.1 % negatively charged residue, predominantly Glu (9.8 %) in the BDR, which is similar to *Proteobacterial* IHFα and IHFβ. We also observed an enrichment of Lys in both AHR and BDR, while Arg is significantly depleted. *Actinobacterial* M1 group, consisting MtbHU, was found to be highly enriched in Arg in the BDR (16.3 %) compared to *Proteobacterial* and *Firmicutes* HU. In *Actinobacteria*, we observed the two separate clades showing distinctive amino acid compositional differences, which was not reported earlier. Their percentages of positively charged residues, small amino acid, aromatic residues and Arg content in the BDR are significantly different. The Actinobacterial M2 group was depleted of aromatic residues In the BDR, and many conserved Phe residues are mutated to Leu, which could introduce higher flexibility in between the two protomers. A comparative structural analysis of the salt-bridge network also suggests a few similarities between HU from *Firmicutes* and *Actinobacteria* (M1 group) and IHF like proteins. We speculate that this could be due to the absence of any other paralogs of HU/IHF proteins in most of the *Firmicutes* and *Actinobacteria*.

## Dimer stabilization differs in HU and IHF

HU/IHF family proteins are obligate dimers and their interface is primarily stabilized by conserved aromatic residues at positions 78, 81 and 117 in MSA1, formed by Phe residues from both the chains. Our study noted that proteins in the HU clade have a single aromatic cluster stabilizing the dimer interface (Fig. 4a), consisting mainly of three Phe residues from each protomer. In IHF clade proteins, an additional aromatic cluster consisting two Phe residue from each protomer was found to be located in the second helix, giving it an extra stabilization and rigidity to the interface (Fig. 4b). In *E. coli* HUαβ structure (PDB: 3O97), the aromatic cluster is formed by F47, F50 and F79 from each protomer while in *Mtb*HU, an extra Phe residue F85 (from each protomer) joins the primary aromatic cluster. F50 and F79 of opposite chains interact via Pi-Pi interaction while the rest of the Phe residues interact through a network of hydrophobic interactions with the surrounding Leu and Ile. While F47, F50, and F79 are highly conserved in all HU/IHF family proteins, F85 is found to be conserved only in *Actinobacterial* M1 group.

**Figure 4.**
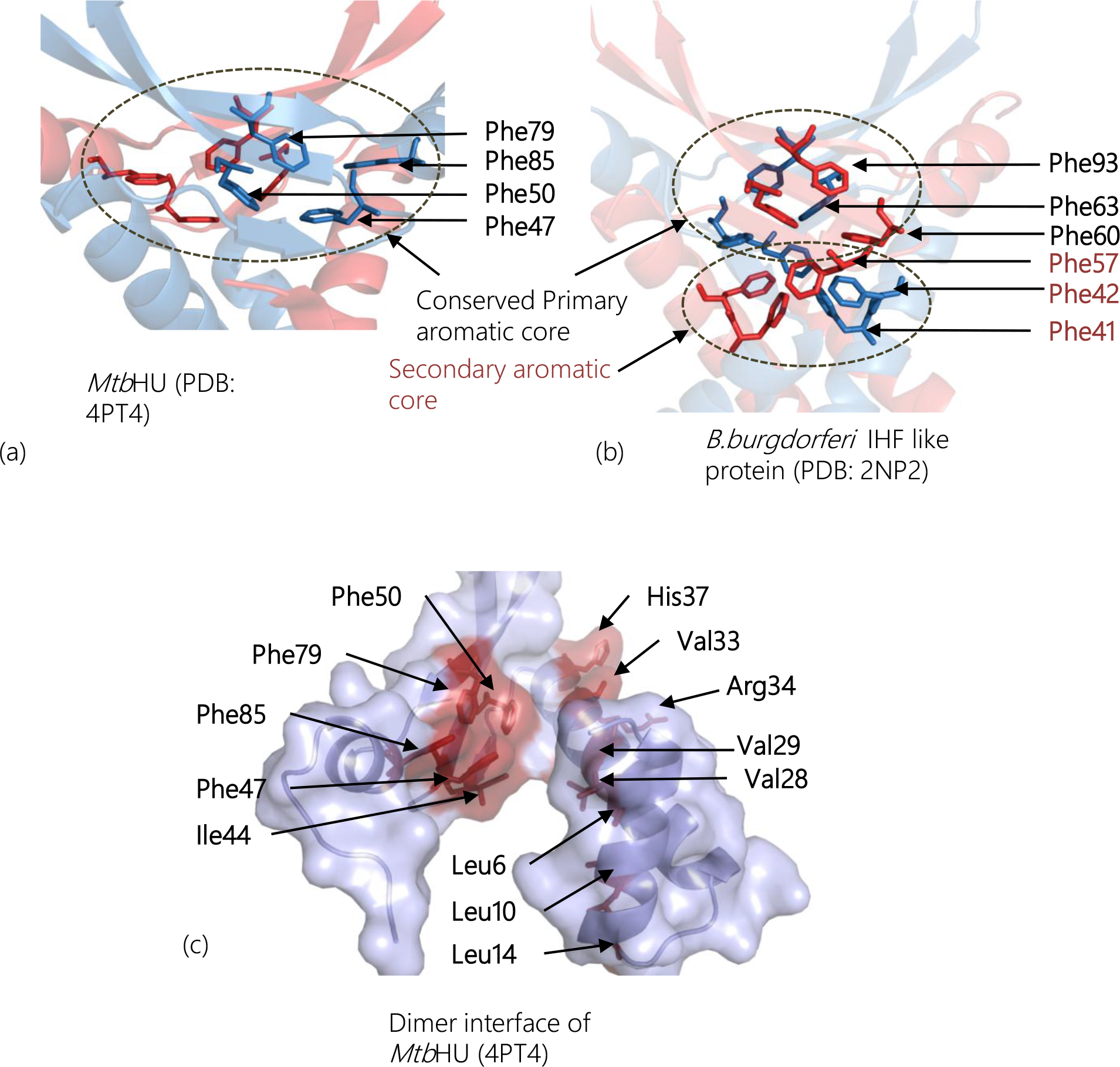
Aromatic core differences in HU and IHFs and dimer interface Hot-Spots. **(a)**Conserved aromatic residues Phe79, Phe50, and Phe47, along with Actinobacteria-specific (M1 subgroup) Phe85 in MtbHU form the primary aromatic core responsible for dimer interface stabilization in MtbHU (4PT4). **(b)** IHF like proteins forms a secondary aromatic core from residues from majorly alpha helical region Phe57, Phe42 and Phe41, which along with the conserved primary aromatic core, stabilizes the dimer interface in B.burgdorferi Hbb(2NP2). **(c)** The dimer interface is majorly formed by aromatic residues in the core, forming Hot-Spots (thermodynamically crucial residues for stabilization of interface, shown in red color surface) and hydrophobic residues majorly Leu and Val from α_1_ and α_2_ helices.

In the *E. coli* IHF, F49, F52 and F81 (from chain A) and F48, F51 and F80 contribute to the primary aromatic core, while F30 and F31 from chain A form a hydrophobic core with M29 and L30 of chain B. We observed that *B.burgdorferi* IHF like protein (PDB: 2NP2) forms a secondary core with F41 and F42 from each protomer (Fig. 4b). This “secondary aromatic core” is a discriminatory difference between HU and IHFs. We also observed an higher occurrence of aromatic residues in the AHR in IHF clades over HU (5.7% and 4.9% in IHFα and IHFβ respectively compared to 2.6% in HU) (Table 1). Computational Ala scanning analysis reveals these aromatic cluster forming residues to be dimer stabilization hot-spots (residues whose alanine mutation destabilizes the interface) (Fig. 4c). Other than aromatic interaction stabilizing the dimer interface, hydrophobic and polar interactions also play a crucial role in its stability and flexibility. In IHF clade proteins, we observed an enrichment of charged residues in the AHR, which “stitches” the two chains with salt bridge and hydrogen bonding interactions. Among the charged residues, Arg and Glu dominate, which contributes to the stabilization of the interface with salt bridges. The differences in the aromatic core taken together with the compositional bias between HU and IHF in the AHR strongly points towards a more rigid dimer interface in IHF compared to HU, which can also influence the overall DNA binding and stabilization as probed in many NMR studies (Wojtuszewski K and Mukerji, 2004).

## Conclusion

Promiscuity or multi-specificity in transcription factors comes with the adaptive flexibility of its binding site upon substrate binding (Nobeli et al., 2009, Fornili et al., 2013). The present report addresses several questions raised during the studies of nucleoid-associated proteins HU and IHF concerning their specificity, DNA bending angle and binding site size. Our analysis provides evolutionary and structural rationalizations of these questions while also determining many new aspects, which affect the specificity and dimer stabilization. This study provides a model example to understand the differences in specificity of proteins belonging to same fold.

In our analysis, we found three major factors, which facilitate the evolution of HU/IHF proteins’ DNA binding multi-specificity. The first factor is the amino acid composition in the DNA-binding region of the protein, which influences the salt bridge constraints, promoting specificity. IHF like proteins contain a higher percentage of both positively and negatively charged residues, which constrains its DNA binding by salt-bridge formation, making it more specific towards its cognate sequence as compared to HU. We have also determined specific positions in HU and IHF clades, which serve as a discriminatory residue positions to classify them. Some class-specific residues like K45 and R76 of *E. coli* IHFα and R42, E44, R46, H54, K75 and H79 of IHFβ (with reference to *E. coli)* promote higher specificity in the DNA binding of IHF compared to its HU homolog. Also, K24 and K45 in IHFα and K27 and R42 in IHFβ extends the positively charged patch on the lateral DNA draping side, which stabilizes the DNA bend and extends the DNA site size in IHF clade proteins compared to HU. The second factor, which influences DNA binding specificity, is conformational flexibility of the binding site due to enrichment of Ala near DNA binding residues. It allows conformational flexibility to the DNA binding side chains in its vicinity. In HU clade proteins, we observed a higher percentage of Ala than IHFs, thus allowing higher side chain conformational flexibility of neighboring DNA binding residues. The third-factor influencing specificity is the type of DNA binding residue. We observed that Arg interacts with DNA bases more predominantly than Lys (Luscombe et al., 2001). Thus, a bias in Arg to Lys ratio influences the DNA binding specificity. We found IHF enriched in Arg while HU in Lys. As HU and IHF are obligate dimers, in effect, each point mutation is duplicated in the DNA binding site and dimerization interface.

Other than the determinants of DNA binding specificity, this report also found an additional aromatic core in the IHF clade proteins, which strengthens the dimer interface and might influence the flexibility in the DNA binding region. Our analysis thus provides a comprehensive evolutionary picture, which explains the origin of multi-specificity using sequence, and structure based DNA binding determinants in HU/IHF fold proteins. The above conclusion regarding determinants of specificity and evolution of promiscuous binding features also holds true for other nucleic acid binding proteins. Thus, this study presents a combined phylogenetic and structural approach, which can be applied to other nucleic acid binding proteins.

## Acknowledgements

D.D., V.N.,and S.R. conducted the studies and wrote the manuscript. The authors thank the members of the Ramakumar’s Lab, Soumitra Ghosh of VN’s Lab, Narmada Sambaturu and Debleena Dey for discussion. This work was supported by a Council of Scientific & Industrial Research (CSIR), India project grant. D.D. thanks, University Grants Commission (UGC), Govt. of India for UGC BSR meritorious fellowship. S.R. thanks, University Grants Commission (UGC), Govt. of India for Emeritus fellowship. The work in V.N. lab is supported by Department of Biotechnology (DBT), Govt. of India.

## Materials and method

### Sequence identification and analysis

Members of HU/IHF family proteins were identified by InterPro ID IPR000119, which represents bacterial histone-like proteins and the dataset of 1112 proteins, were created. It excludes HU/IHF family proteins from phylum *Bacteriodetes* and *Eukaryotes* due to high sequence divergence and poor multiple sequence alignment. The dataset was further validated by PSI-BLAST runs performed on experimentally confirmed HU and IHF sequences as aninitial query against the non-redundant sequence database (Table S1). CD-HIT server (Li and Godzik 2006) was used to remove redundancy in sequence data. Two steps were used to reduce redundancy in the sequence dataset. Firstly, a cutoff of 90% sequence identity was used to remove redundancy in the phyla. The sequences were aligned using CLUSTALW in MEGA 6.0 (Kumar et al., 2008). The aligned block, corresponding to the fold representing HU/IHF family proteins, was extracted (N-terminal M1 to C-terminal S90 to reference sequence HU *Mycobacterium tuberculosis*). The representative sequences from different phyla were pooled and redundancy was removed with a 70% sequence identity cutoff. Phyla specific HU datasets were prepared with a redundancy cutoff of 90% sequence identity.

### Phylogenetic analysis and information theoretic methods

Evolutionary trace method partitions the dendrogram and assorts multiple sequence alignment (MSA) positions into invariant, class specific and variable types. Each partition or “trace” consist its cluster (class) specific positions. We used an evolutionary trace (ET) method based on Kitsch algorithm to build a rooted phylogenetic tree (Innis et al., 2000), using Trace Suite II server. Also to confirm the topology of the phylogenetic tree, neighbor-joining (NJ) tree and maximum likelihood tree were prepared. ProtTest 1.4 (Abascal et al., 2005) was used to determine the best-fit amino acid substitution model and parameter from values for each data set. In each case, the (Le and Gascuel 2008) LG model was the best fit according to the Akaike information criterion. PHYML 3.0 server was used to run the ML analysis. The Subtree Pruning and Regrafting method were used to search tree topology. Branch support for the resulting topology was determined by the Shimodaria-Hasegawa-like approximate likelihood ratio test.

BioEdit was used to calculate the position specific Shannon’s entropy and residue composition for MSA (Hall 1999).Analysis of positional conservation cannot be done without taking into account the conservation of essential nature of the side chain. Thus, amino acid groups can be defined for which the group specific Shannon’s entropy can be calculated (Guharoy and Chakrabarti 2005, Sanchez-Flores 2007). A five class model: Hydrophobic (Gly, Ala, Leu, Ile, Met and Val) Hydrophilic (Pro, Thr, Ser, Cys, Asn and Gln) Aromatic (Tyr, Phe, and Trp), Positive charged (Lys, Arg and His) and Negatively charged (Asp and Glu), served the best for our analysis. Histidine in HU homologs is usually found in the beta sheet DNA binding region, thus, it is grouped along with the positively charged residues than aromatic ones. The normalized Shannon’s entropy is given by the following expression.

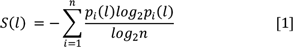

Where *p_i_(l)* represents the frequency of *i* class of residues at position *l* in the multiple sequence alignment and n represents the number of amino acid groups depending the classification criteria. For our calculation *n*=5 (five classes) was chosen. The higher entropy implies higher variability in the given position in multiple sequence alignment and *vice versa*. The normalized class specific entropy for different traces was represented in heatmap format, where red to blue spectrum correspond to avariable to conserved positions respectively. We have considered entropy range 0-0.3 as conserved, 0.31-0.7 as intermediate and 0.71-1 as variable positions. In entropy analysis, positions with “gap” frequency are more than 0.7, is ignored (represented as gray color in entropy heatmap). To calculate statistical significance in amino acid composition, we used the web server Composition profiler (http://www.cprofiler.org/), in which we used one set of amino acid composition (of a clade) as background while the other as a query (Vacic et al., 2007) (Supplementary Table S6-9). Geneious software was used to obtain consensus sequence, motif logo, and MSA visualization. Sequence divergence parameters from MSA were calculated using ALISTAT server (http://caps.ncbs.res.in/iws/alistat_ali.html). For Homology modeling of *E. coli* HU chains, PHYRE server was used.

### Structural analysis

Interactions were classified into three types a) Intra-chain b) inter-chain (for dimer interface) and c) DNA-protein. All the interactions were calculated using Accelrys Discovery studio interaction calculator. Robetta alanine scanning (Kortemme et al., 2004) was used to predict hot-spots at the dimer interface. Hot-spot residues can be defined as those positions for which alanine mutations have destabilizing effects on ΔΔG_bind_ of more than 1 kcal/mol. All the Figures were generated using PyMOL (deLano Scientific). Hydrophobic interactions are classified into aromatic interactions (Pi-Pi, Pi-Pi T-shaped and Pi-cation) and non aromatic hydrophobic interaction. For the latter case 8Å cutoff was taken (Tina et al., 2007), while aromatic interaction used standard geometric parameters.

